# LL-37 and Citrullinated-LL-37 elicited Transcriptome and Cytokine Profiles in Human Bronchial Epithelial Cells: Citrullination dampens the inflammatory biosignature

**DOI:** 10.64898/2026.09.20.753030

**Authors:** Padmanie Ramotar, Dina H D Mostafa, Mahadevappa Hemshekhar, Courtney L Marshall, Christopher D Pascoe, Neeloffer Mookherjee

## Abstract

The human host defence peptide LL-37 mediates pleiotropic immunomodulatory functions that can be both pro- and anti-inflammatory. Under inflammatory conditions in the lungs, LL-37 is susceptible to the post-translational modification (PTM) citrullination. The impact of this PTM on LL-37’s immunomodulatory functions is not fully understood. Therefore, we characterized the transcriptome and cytokine profile in response to LL-37 and citrullinated-LL-37 (citLL-37), in human bronchial epithelial cells (HBEC). Cells were stimulated with either LL-37 or citLL-37 (0.5 μM), or a scrambled peptide (sLL-37). RNA after 4 hours (h) was analyzed with NanoString nCounter Host Response Panel, and 96 cytokines were examined (after 24 h) in tissue culture supernatants (Luminex platform). Genes and cytokines with a Log2 fold-change ≥0.5 with *p*<0.05 compared to unstimulated cells were considered differentially expressed (DE). These studies revealed overlapping and distinct biosignatures; There were 118 DE genes with LL-37 and 71 with citLL-37, of which magnitude of change was significantly different for 32 genes. 16 out of 17 DE genes altered by both peptides were directly related to inflammation, and these were significantly less (by ∼45%) with citLL-37 compared to LL-37. Similarly, inflammatory cytokines enhanced in response to LL-37 were significantly less (by 37-94%) with citLL-37. Overall, our findings indicate that citrullination of LL-37 does not abrogate cellular response in HBEC, instead dampens the inflammatory biosignature induced by LL-37.

## INTRODUCTION

Cationic host defence peptides (CHDPs) are small amphipathic peptides found in all living organisms. Cathelicidins are one of the most well characterized family of CHDPs in mammals [1]. The sole human cathelicidin peptide is LL-37, which is derived from hCAP18 encoded by the cathelicidin gene *CAMP*. LL-37 exhibits wide spectrum antimicrobial activity against bacteria, viruses and fungi [2,3]. Moreover, immunity-related functions of LL-37 play a critical role in the resolution of infections, regulation of inflammation and epithelial wound repair [1,4–6]. In the context of its immunomodulatory functions, LL-37 can elicit opposing functions as it facilitates both pro- and anti-inflammatory responses depending on the microenvironment such as type of extracellular factors and external stimuli and concentration of the biologically active mature peptide [1,4,7]. Likewise, in the lungs, LL-37 can both promote and suppress inflammation; LL-37 can promote airway inflammation [8–10], and facilitate recruitment of inflammatory leukocytes to the lungs [9,11]. In contrast, LL-37 can be protective by decreasing inflammatory responses in models of acute lung injury [12,13], and shown to resolve airway inflammation in various *in vitro* and *in vivo* models of respiratory disease [14–17]. Although the role of LL-37 in regulating inflammation is well appreciated, the underlying mechanisms that coordinate the shift between the pro- and anti-inflammatory effects of the peptide have yet to be completely resolved.

LL-37 is susceptible to citrullination which is a post-translational modification (PTM) where positively charged arginine residues are converted to neutral citrulline typically catalyzed by peptidyl-arginine deaminase PAD enzymes [18]. PAD enzymes are elevated in the lungs during airway inflammation [19], and citrullinated-LL-37 (citLL-37) is detected in the bronchoalveolar lavage fluid (BALF) of humans, thus LL-37 can get citrullinated under inflammatory conditions in the lungs [18]. This is corroborated by a recent study in human bronchial epithelial cells (HBEC) demonstrating an increase in PAD enzyme and citrullination levels during human rhinovirus infection, leading to the citrullination of LL-37 and consequent impairment of the peptide-mediated antiviral function [20]. Although citrullination compromises LL-37’s antimicrobial functions [18,20,21], the impact of this PTM on the immunomodulatory functions of the peptide remains largely unresolved.

We have previously shown that citrullination does not abrogate LL-37’s immunomodulatory functions in HBECs [9,14]. Therefore, in this study, we profiled LL-37- and citLL-37-mediated changes in HBEC transcriptome and secreted cytokines to define the impact of citrullination on the peptide-induced responses in airway epithelium. The findings of this study provide the foundational dataset to detail the differences and commonalities between LL-37- and citLL-37-mediated responses in HBEC. We show concordance between the transcriptional and cytokine responses elicited by these peptides, providing protein-level confirmation of selected gene expression and reinforcing the robustness of the transcriptomic dataset reported in this study. Overall, this study demonstrates that citrullination of LL-37 does not abrogate its biological responses in HBEC, instead it markedly attenuates the inflammatory molecular signature induced by the peptide. This study provides compelling evidence to define alteration of LL-37-driven epithelial responses following citrullination and the framework for future mechanistic studies to delineate the role of PTMs in the functions of LL-37 and other CHDPs in the lungs.

## RESULTS

### LL-37 and citLL-37 mediate overlapping and distinct transcriptional changes in human bronchial epithelial cells

HBEC3-KT cells (ATCC CRL-4051™) were treated with either 0.50 µM of LL-37, citLL-37, or a scrambled peptide (sLL-37) for 4 hours (h), and transcriptional responses were examined using the NanoString™ nCounter® Host Response Panel. Principal component analysis (PCA) was conducted on the delta expression values after subtracting the expression values obtained from unstimulated cells. The first two principal components (PC1 and PC2) represented 30.8% and 18.8% of the total variance, respectively (Figure 1). The samples separated primarily along the PC1, with LL-37-treated cells clustering on the negative axis and sLL-37 treated cells clustering on the positive axis. citLL-37-treated cells clustered between LL-37 and sLL-37. All three peptide-treated samples separated along the PC1 axis. These results suggested that the transcriptional profile elicited by each peptide treatment was overall distinct from each other.

**Figure 1:**
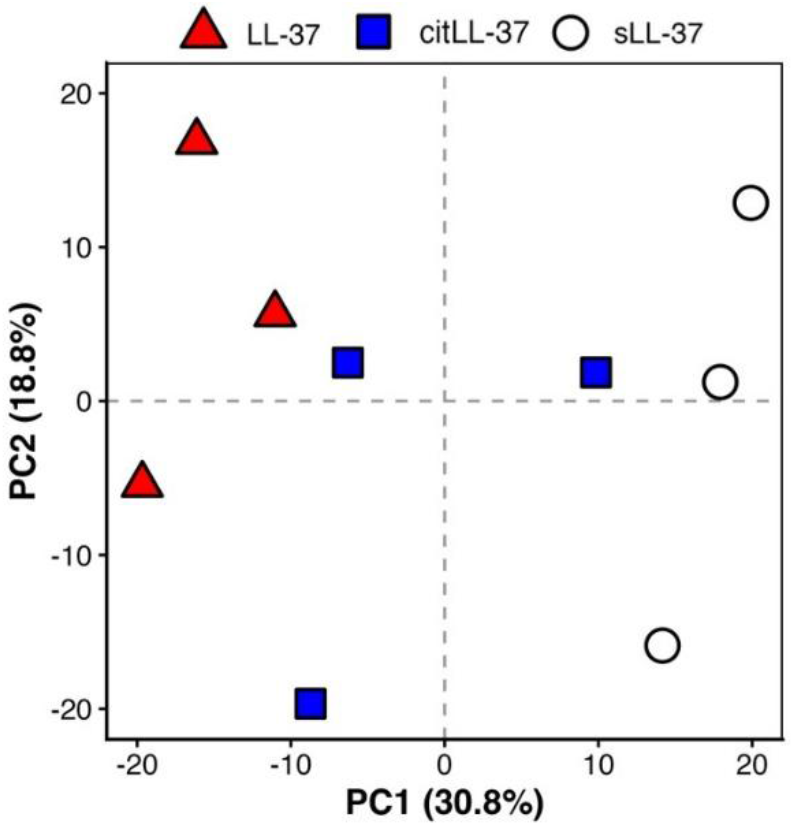
Principal Component Analysis (PCA) of LL-37, citLL-37 and sLL-37 mediated changes in the human bronchial epithelial cell transcriptome. HBEC3-KT cells were stimulated with LL-37, citLL-37, or sLL-37 (0.50 µM) and RNA (n=3) collected after 4 h was analyzed using the NanoString nCounter® Host Response Panel. Principal component analysis (PCA) of each condition after background subtraction of expression in unstimulated cells for each gene.

Differential gene expression analysis was performed using the limma package with empirical Bayes moderation. Genes with a Log2 fold-change ≥ 0.5 and a *p*-value < 0.05, compared to unstimulated cells, were considered as significantly differentially expressed (DE). As sLL-37 does not mediate immunomodulatory functions like LL-37 and is used as a negative peptide [9,14], 22 DE genes (10 up-regulated and 12 down-regulated) with sLL-37 were removed from further analyses (Supplementary Table 1). The resultant gene expression matrix identified 118 DE genes with LL-37 (Figure 2A and Supplementary Table 2) and 71 DE genes with citLL-37 (Figure 2B and Supplementary Table 2). These results demonstrated that citrullination of LL-37 does not abrogate but alters the transcriptional response, with less DE genes compared to that mediated by LL-37 in HBECs.

**Figure 2:**
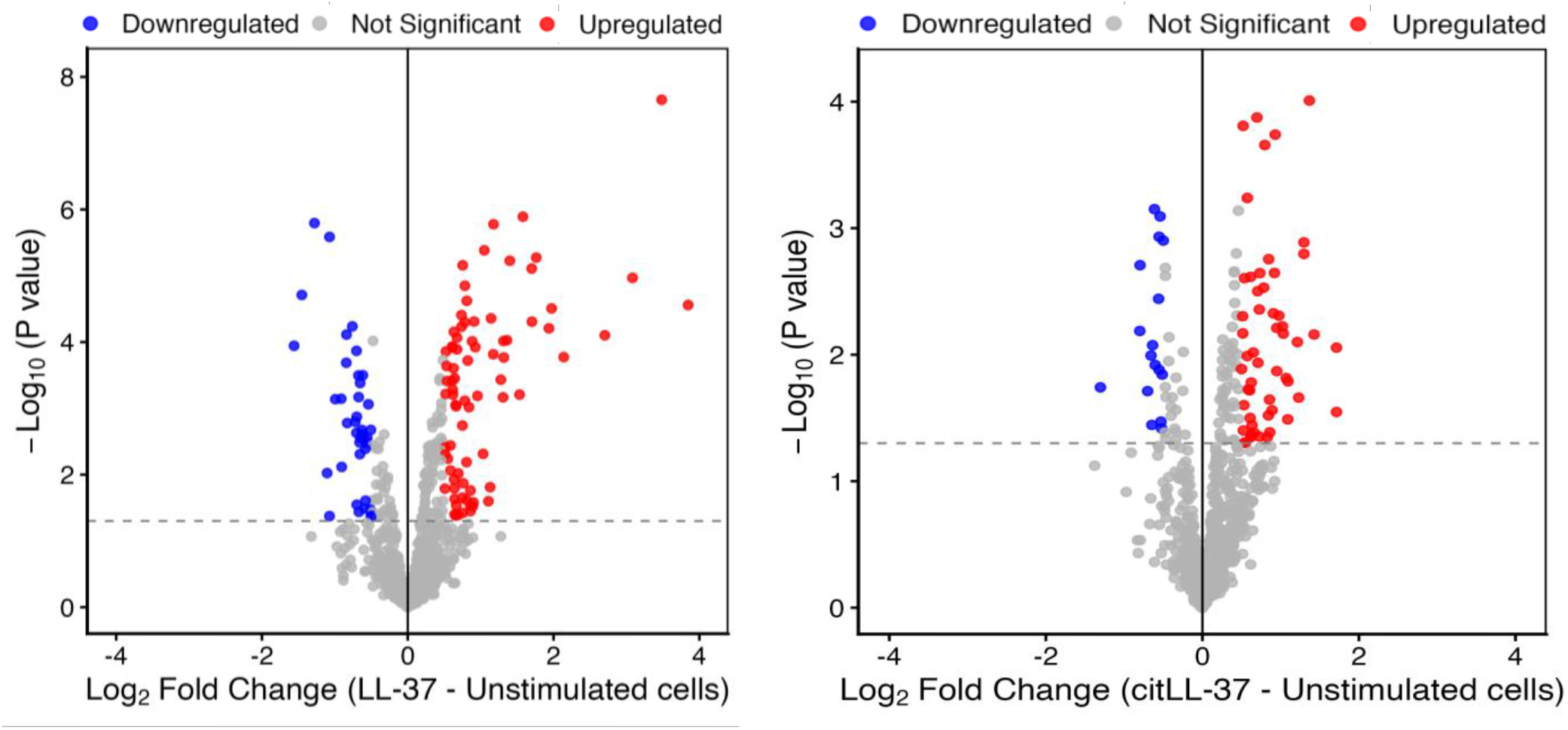
Transcriptional changes mediated by LL-37 and citLL-37 in human bronchial epithelial cells. HBEC3-KT cells were stimulated with **(A)** LL-37 or **(B)** citLL-37, and RNA collected after 4 h was analyzed using the NanoString nCounter® Host Response Panel (n=3). Volcano plots showing the differentially expressed genes for each peptide treatment vs unstimulated cells. The X axis represents Log2 fold-change of peptide-mediated transcriptional response compared to unstimulated cells. The Y axis represents -Log10 (*p*-value). Red dots indicate significant increase, blue dot indicate significant decrease, and grey indicates non-significant change, relative to unstimulated cells.

Comparative analyses of DE genes in response to either LL-37 or citLL-37 revealed 32 DE genes with significant quantitative differences in magnitude of change in response to citLL-37 compared to LL-37 (Table 1). Of these, 11 DE genes were uniquely altered in response to LL-37 (4 increased, 7 decreased), and 4 DE genes (3 increased, 1 decreased) were unique to citLL-37 (Table 1). There were 17 genes that were common between LL-37- and citLL-37-mediated DE genes. Of these 17 DE genes, 16 were directly related to inflammation (*PTGS2, CXCL8, CSF2, IL1A, IL1RL1, VEGFA, CXCL3, CXCL1, SOCS3, IL1B, PELI1, PLAUR, IRF1, TXNIP, MCL1 and TRIM6*), and the magnitude of change for all of these was significantly less with citLL-37 (∼45%) compared to LL-37 (Table 1).

**Table 1:** LL-37 and citLL-37 DE genes with significant difference in magnitude of response.

| Gene | LL-37 |  | citLL-37 |  | LL-37 – citLL-37 |  |
| --- | --- | --- | --- | --- | --- | --- |
|  | Log2 Fold-Change* | p-value | Log2 Fold-Change* | p-value | Log2 Fold-Change | p-value |
| *PTGS2* | 3.85 | 2.76E-05 | 1.71 | 8.79E-03 | 2.13 | 2.37E-03 |
| *CXCL8* | 3.48 | 2.21E-08 | 1.37 | 9.80E-05 | 2.12 | 2.29E-06 |
| *CSF2* | 3.08 | 1.07E-05 | 1.09 | 1.62E-02 | 1.99 | 3.81E-04 |
| *HMOX1* | 2.70 | 7.89E-05 | 1.43 | 6.92E-03 | 1.28 | 1.26E-02 |
| *IL1A* | 1.70 | 7.78E-06 | 0.53 | 2.50E-02 | 1.17 | 1.76E-04 |
| *IL1RL1* | 2.14 | 1.68E-04 | 1.07 | 1.53E-02 | 1.07 | 1.53E-02 |
| *VEGFA* | 1.97 | 3.10E-05 | 0.95 | 6.14E-03 | 1.02 | 3.95E-03 |
| *CXCL3* | 1.58 | 1.28E-06 | 0.62 | 2.42E-03 | 0.96 | 9.29E-05 |
| *CXCL1* | 1.76 | 5.31E-06 | 0.85 | 1.76E-03 | 0.92 | 1.02E-03 |
| *SOCS3* | 1.70 | 4.92E-05 | 0.91 | 4.71E-03 | 0.80 | 9.81E-03 |
| *IL1B* | 1.36 | 9.37E-05 | 0.60 | 1.91E-02 | 0.76 | 5.77E-03 |
| *PELI1* | 1.31 | 9.63E-05 | 0.59 | 1.88E-02 | 0.73 | 6.07E-03 |
| *PLAUR* | 1.31 | 6.81E-04 | 0.62 | 4.49E-02 | 0.69 | 2.82E-02 |
| *IRF1* | -1.28 | 1.60E-06 | -0.61 | 7.07E-04 | -0.67 | 3.84E-04 |
| *TXNIP* | -1.45 | 1.95E-05 | -0.80 | 1.96E-03 | -0.66 | 6.45E-03 |
| *MCL1* | 1.14 | 4.39E-05 | 0.50 | 1.30E-02 | 0.64 | 3.08E-03 |
| *TRIM6* | -1.07 | 2.59E-06 | -0.50 | 1.25E-03 | -0.57 | 4.72E-04 |
| *KIR3DL1/2* | -0.15 | 6.40E-01 | 0.86 | 2.26E-02 | -1.01 | 9.96E-03 |
| *FOS* | 0.09 | 6.97E-01 | -0.80 | 6.49E-03 | 0.89 | 3.36E-03 |
| *AIF1* | -0.02 | 9.35E-01 | 0.78 | 2.95E-03 | -0.80 | 2.58E-03 |
| *RSAD2* | -0.08 | 7.52E-01 | 0.55 | 4.99E-02 | -0.63 | 2.87E-02 |
| *IL17C* | -1.11 | 9.46E-03 | 0.51 | 1.70E-01 | -1.62 | 8.82E-04 |
| *IL24* | 1.53 | 6.20E-04 | 0.36 | 2.74E-01 | 1.17 | 3.73E-03 |
| *CXCL2* | 1.28 | 3.69E-04 | 0.23 | 3.57E-01 | 1.04 | 1.56E-03 |
| *RNASEL* | -1.56 | 1.14E-04 | -0.55 | 5.44E-02 | -1.01 | 2.71E-03 |
| *CD274* | 1.18 | 1.67E-06 | 0.39 | 8.09E-03 | 0.79 | 5.36E-05 |
| *RNF135* | -0.91 | 7.16E-04 | -0.28 | 1.70E-01 | -0.63 | 7.49E-03 |
| *NFATC3* | -0.65 | 2.76E-03 | -0.06 | 7.23E-01 | -0.59 | 5.01E-03 |
| *KDM6B* | 0.96 | 6.48E-04 | 0.38 | 8.30E-02 | 0.58 | 1.42E-02 |
| *OASL* | -0.59 | 3.24E-02 | -0.06 | 8.15E-01 | -0.53 | 4.88E-02 |
| *TNF* | -0.99 | 7.26E-04 | -0.47 | 4.60E-02 | -0.52 | 2.96E-02 |
| *TNFSF10* | -0.84 | 7.71E-05 | -0.34 | 2.73E-02 | -0.50 | 3.19E-03 |

\*Compared to unstimulated cells. Grey = genes not DE and/or not significant.

KEGG pathway enrichment analysis demonstrated that the top 4 enriched pathways associated with the 32 DE genes with significant quantitative differences in magnitude between LL-37 and citLL-37 (Table 1) were all related to inflammatory processes (Table 2). These pathways primarily constituted of DE genes that were significantly less in response to citLL-37 compared to LL-37. These pathways represented key inflammatory mediators TNF, IL-17 and NF-κB signaling, and related to chronic inflammatory disease rheumatoid arthritis. Together, these findings demonstrated that citrullination of LL-37 does not abrogate the peptide-mediated transcriptional responses, instead dampens the LL-37-induced inflammatory biosignature.

**Table 2:** Pathway enrichment from LL-37 and citLL-37 DE genes with differential magnitude of response.

| Pathways | Gene Count | Adjusted *p*-value | Genes |
| --- | --- | --- | --- |
| Rheumatoid arthritis | 10 | 4.62E-04 | *CXCL8, CXCL3, IL1A, CSF2, CXCL1, CXCL2, FOS, VEGFA, IL1B, TNF* |
| IL-17 signaling pathway | 10 | 1.40E-03 | *CXCL8, CXCL3, CSF2, IL17C, CXCL1, CXCL2, PTGS2, FOS, IL1B, TNF* |
| TNF signaling pathway | 10 | 3.48E-03 | *CXCL3, CSF2, IRF1, CXCL1, CXCL2, PTGS2, FOS, IL1B, SOCS3, TNF* |
| NF-κB signaling pathway | 7 | 4.99E-02 | *CXCL8, CXCL3, CXCL1, CXCL2, PTGS2, IL1B, TNF* |

### Cytokines induced in response to LL-37 and citLL-37 in human bronchial epithelial cells

As cytokines and chemokines were primarily represented in the top 4 enriched pathways identified from the transcriptomic analysis (Table 2), we further profiled the abundance of 96 cytokines secreted from HBEC3-KT cells following stimulation with either LL-37, citLL-37 or sLL-37 (0.5 µM each). Tissue culture (TC) supernatants collected after 24 h were examined for protein abundance of 96 human cytokines and chemokines using a multiplex assay. Stimulation with sLL-37 resulted in the enhancement of 5 cytokines (MIP-3α, sCD40L, IL-11, IL-20 and CXCL16) and these were removed from the analysis. The abundance of 15 cytokines was significantly enhanced by either LL-37 and/or citLL-37 (Table 3). Of these, 14 cytokines were significantly increased in response to LL-37, and 6 in response to citLL-37, with 5 cytokines significantly enhanced by both peptides (Table 3). Similar to the transcriptional biosignature, LL-37 induced a broader inflammatory response compared to citLL-37 (Supplementary Table 3). Citrullination of LL-37 significantly decreased LL-37-induced abundance of inflammatory mediators such as GM-CSF, IL-8, I-TAC, IL-6, IP-10, Fractalkine and Eotaxin between 37-94% (Supplementary Table 3). These results suggested that citrullination attenuates the pro-inflammatory cytokine response elicited by LL-37.

**Table 3.** Cytokines and chemokines significantly altered by LL-37 and/or citLL-37.

| Protein | LL-37 |  | citLL-37 |  |
| --- | --- | --- | --- | --- |
|  | Log2 Fold-Change* | *p*-value | Log2 Fold-Change* | *p*-value |
| GM-CSF | 4.33 | 3.50E-10 | 0.27 | 3.03E-01 |
| I-TAC | 0.65 | 2.76E-07 | -0.02 | 7.46E-01 |
| IL-6 | 1.02 | 2.09E-06 | 0.25 | 6.53E-02 |
| IL-8 | 4.27 | 4.24E-06 | 1.47 | 2.08E-02 |
| IP-10 | 2.89 | 7.35E-06 | 0.93 | 3.68E-02 |
| Fractalkine | 2.87 | 1.66E-04 | 0.20 | 7.18E-01 |
| GROα | 2.35 | 3.95E-04 | 1.57 | 7.22E-03 |
| TNFα | 1.25 | 6.45E-04 | 0.72 | 2.32E-02 |
| Eotaxin | 1.25 | 1.18E-03 | 0.46 | 1.53E-01 |
| IL-1α | 2.07 | 3.03E-03 | 0.85 | 1.57E-01 |
| IFNγ | 0.68 | 4.96E-03 | 0.20 | 3.38E-01 |
| RANTES | 0.65 | 6.03E-03 | 0.55 | 1.68E-02 |
| G-CSF | 0.80 | 2.04E-02 | 0.22 | 4.73E-01 |
| TNFβ | 1.14 | 3.57E-02 | 0.41 | 4.11E-01 |
| SCF | -1.46 | 3.58E-02 | 1.38 | 4.55E-02 |

\*Compared to unstimulated cells. Grey denotes cytokines not significantly changed.

### Concordance of transcriptional and cytokine responses induced by LL-37 and citLL-37

To identify cytokine targets that were changed at both the transcript and protein level by the peptides, the expression matrix obtained from the NanoString analysis (Supplementary Table 2) was compared to dataset obtained from cytokine profile (Supplementary Table 3). There were 17 candidates that were detected in both the transcriptomic and cytokine datasets (Supplementary Table 4). Of these, there were 6 cytokines with concordant change in response at both transcriptional and protein levels (Table 4). LL-37 enhanced the abundance of *CXCL8* (IL-8), *CSF2* (GM-CSF), *CXCL1* (GROα), *IL1A* (IL-1A) and *CXCL11* (ITAC), both at the gene and protein level, all of which play a role in inflammation. Protein abundance of TNFα was also significantly enhanced by LL-37 (Table 4). All these inflammatory candidates were significantly less (by ∼50%) in response to citLL-37 (Table 4). These findings identified specific inflammatory cytokines enhanced by LL-37, with concordance between transcriptional and protein response, and demonstrated significant dampening of these inflammatory responses by citrullination of the peptide.

**Table 4:**
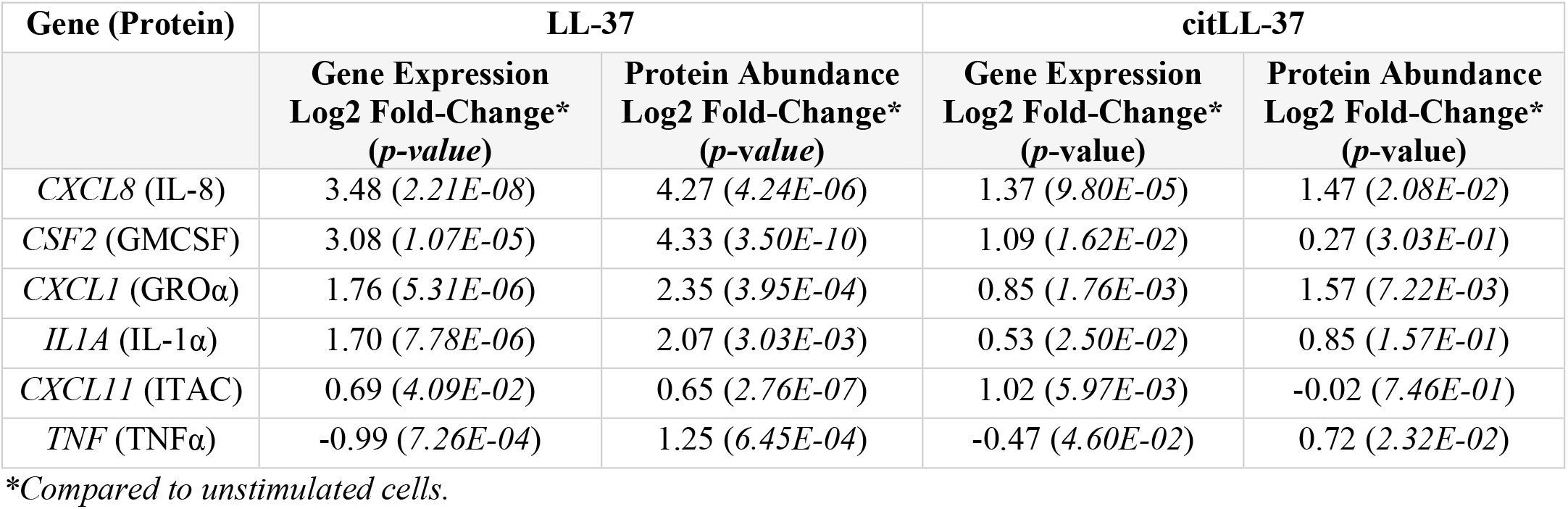
Concordant transcriptional and cytokine response elicited by LL-37 and citLL-37.

| Gene (Protein) | LL-37 |  | citLL-37 |  |
| --- | --- | --- | --- | --- |
|  | Gene Expression<br>Log2 Fold-Change*<br>(*p*-value) | Protein Abundance<br>Log2 Fold-Change*<br>(*p*-value) | Gene Expression<br>Log2 Fold-Change*<br>(*p*-value) | Protein Abundance<br>Log2 Fold-Change*<br>(*p*-value) |
| *CXCL8* (IL-8) | 3.48 (2.21E-08) | 4.27 (4.24E-06) | 1.37 (9.80E-05) | 1.47 (2.08E-02) |
| *CSF2* (GMCSF) | 3.08 (1.07E-05) | 4.33 (3.50E-10) | 1.09 (1.62E-02) | 0.27 (3.03E-01) |
| *CXCL1* (GROα) | 1.76 (5.31E-06) | 2.35 (3.95E-04) | 0.85 (1.76E-03) | 1.57 (7.22E-03) |
| *IL1A* (IL-1α) | 1.70 (7.78E-06) | 2.07 (3.03E-03) | 0.53 (2.50E-02) | 0.85 (1.57E-01) |
| *CXCL11* (ITAC) | 0.69 (4.09E-02) | 0.65 (2.76E-07) | 1.02 (5.97E-03) | -0.02 (7.46E-01) |
| *TNF* (TNFα) | -0.99 (7.26E-04) | 1.25 (6.45E-04) | -0.47 (4.60E-02) | 0.72 (2.32E-02) |

\*Compared to unstimulated cells.

## Discussion

In this study, we characterized the transcriptional and cytokine response profiles elicited by physiologically relevant concentrations of LL-37 and citLL-37 [9,22]. Citrullination is an irreversible PTM that has garnered interest due to its role in influencing biological processes related to immune function, inflammation and regulation of gene expression [23]. To our knowledge, this is the first study to provide a comparative analysis of LL-37- and citLL-37-mediated responses in HBEC transcriptome and cytokine profiles. Overall, the findings reported in this study suggest that citrullination does not abrogate LL-37-induced cellular responses, instead it dampens LL-37-mediated inflammatory signature in the bronchial epithelia.

Citrullinated forms of LL-37 are detected in the human lungs [18]. During airway inflammation, there is an increase in both PAD enzymes and LL-37, consequently LL-37 is likely to get citrullinated [18,19]. Citrullination of LL-37 suppresses the peptide’s direct antiviral activity against respiratory viruses [20], as well as its antibacterial functions under inflammatory conditions in the airways [19]. The loss of LL-37’s direct antimicrobial function following citrullination is attributed to the loss of positive charge and consequently its binding capacity with microbial membranes [18,20]. However, LL-37 is known to elicit a wide range of immune-related functions which includes enhancing the host’s innate immune responses to facilitate infection resolution, and in contrast negative regulation of inflammation and maintenance of immune homeostasis [1,24–26]. Limited studies have demonstrated that citrullination suppresses some of the immunomodulatory activities of LL-37 by reducing its ability to bind to bacterial LPS and regulate TLR-signaling, as well as interferes in the peptide’s interaction with extracellular DNA [21,27]. This suggests that citrullination could dampen LL-37’s role in facilitating host’s response to microbial challenge and shifts the immune response to a less inflammatory phenotype. This is corroborated by our previous study in HBECs demonstrating that citrullination reduces LL-37-mediated COX-2 signaling and downstream pro-inflammatory responses such as prostaglandin PGE2 and chemokines [9], and impairs the peptide’s ability to promote neutrophil migration under inflammatory conditions [14]. Consistent with this, findings reported in this study show that LL-37-induced upregulation of *PTGS2* (gene encoding PGE2) and production of neutrophil-attracting chemokines such as IL-8 are significantly reduced by at least 4-fold in response to citLL-37 compared to the native peptide. Comparative analysis of our transcriptomic and cytokine datasets clearly demonstrates that citrullination of LL-37 shifts the HBEC biosignature to a reduced inflammatory phenotype. Collectively, these studies suggest that citrullination of LL-37 under inflammatory conditions may be a mechanism to negatively regulate inflammation and promote immune homeostasis, a concept that warrants further functional investigation. Notably, results of this study show that LL-37-induced transcriptional and cytokine response in HBEC are not abrogated, instead quantitatively less, in response to the citrullinated form of LL-37. Similarly, using lipidomics, we have previously shown that both LL-37 and citLL-37 induces the production of oxylipins, and that citrullination of LL-37 does not abrogate but changes oxylipin response in HBECs [9]. LL-37-induced pro-inflammatory bioactive lipids like prostaglandins are selectively suppressed by the citrullination of the peptide [9]. Here, we show that citLL-37 shifts the transcriptional gene signature by suppressing the pro-inflammatory responses activated by LL-37. Across the enriched pathways, the dominant biosignature following citrullination of LL-37 reflected attenuation of critical pro-inflammatory cytokines / chemokines associated with TNFα-, IL-17- and NF-κB -signaling pathways, as well as related to chronic inflammatory disease rheumatoid arthritis. Taken together, these studies indicate that citrullination may disrupt LL-37’s ability to effectively engage with canonical pro-inflammatory signaling, underlying mechanisms of which warrants further investigation.

A recent study suggested that changes in the α-helical fold of LL-37 by citrullination results in the loss of the antimicrobial activity, whereas PTMs such as formylation and acetylation that only modifies the N-terminal maintaining α-helicity of LL-37 retains the peptide’s antimicrobial functions [28]. In contrast, PTMs (formylation and acetylation) disrupting the di-leucine motif at the N-terminal of LL-37, but not citrullination, impairs leukocyte responses such as autophagy [28]. Overall, previous studies indicate that citrullination of LL-37 abrogates antimicrobial functions [18–20], but selectively alters the peptide-mediated immunomodulatory responses [9,14,28]. Thus, it is likely that different PTMs differentially change LL-37’s antimicrobial and immunity-related functions. Each PTM can alter the peptide’s charge distribution and α-helical fold, consequently altering engagement of LL-37 with its direct interacting protein partners or receptors in a distinct manner, making it plausible that each PTM alters LL-37-mediated biological activity differently. To that end, a layer of complexity to consider is that LL-37 is known to engage multiple receptors and accessory proteins depending on the cellular microenvironment to mediate its pleiotropic responses [24,29,30]. We have shown that receptors involved in LL-37-mediated pro- and anti-inflammatory responses are different [31]. Therefore, it is likely that differences in immunomodulatory responses mediated by LL-37 and citLL-37 may be due to differential receptor engagement and/or change in affinity to binding with its interacting protein partners. Therefore, the structure-function relationship governing the different post-translationally modified forms of LL-37, including citrullinated peptides, needs to be systematically characterized in future studies.

A major consideration is that the degree of citrullination may differentially affect LL-37’s immune properties. There are five arginine residues in LL-37, and different citrullinated forms of LL-37 are found in the human lungs [18]. In this study, we used citLL-37 where all five arginine were citrullinated. Mitigation of antimicrobial activity of LL-37 by citrullination is compromised with partially citrullinated peptide where 3 arginine residues are modified [19]. As LL-37 with varying numbers of arginine residues citrullinated are found in the airways [18], it will be imperative to compare immunomodulatory functions between LL-37 with its different physiologically relevant citrullinated forms, which may provide insight into mechanisms associated with modulation of inflammatory pathways in respiratory disease. Other limitations of this study include use of an immortalized cell line for HBEC and examination of responses at a single time point. It is possible that citrullination of the peptide changes the kinetics of LL-37-mediated cellular response, which needs further investigation. Additionally, the transcriptome and cytokine profiles detailed in the findings of this study needs to be confirmed in primary cells isolated from human lungs. However, for some of the chemokines reported in this study, responses induced by both LL-37 and citLL-37 are consistent with our previous study using primary human bronchial epithelial cells [14]. Therefore, the datasets detailed in this study provide objective readouts to systematically examine LL-37- and citLL-37-mediated immunomodulatory responses in primary HBECs isolated from the lungs under different conditions and different cohorts of respiratory disease.

Overall, this study broadly characterizes LL-37- and citLL-37-mediated transcriptional and cytokine responses in HBEC. The datasets detailed in this study, along with our previous lipidomics study [9], provides objective endpoints as a valuable resource for researchers to examine how citrullination alters LL-37-mediated biological processes in bronchial epithelia under different pulmonary conditions. Secondly, the findings of this study suggest that citrullination maybe a regulatory biological phenomenon to dampen the inflammatory biosignature triggered by LL-37 and facilitate immune homeostasis.

## Materials and Methods

### Peptides

The cationic peptides LL-37, citLL-37, and sLL-37 (scrambled peptide) were synthesized and purchased from Innovagen AB (Lund, Sweden) and stored in -20°C until use (Table 5). Peptides were reconstituted in endotoxin-free water (Fisher Scientific, Cat# SH3052901), aliquoted and stored in glass vials at -20°C for a maximum of 3 months. Prior to use, a reconstituted peptide aliquot was thawed at room temperature (RT), sonicated for 30 seconds in a water bath and vortexed for 15 seconds.

**Table 5:**
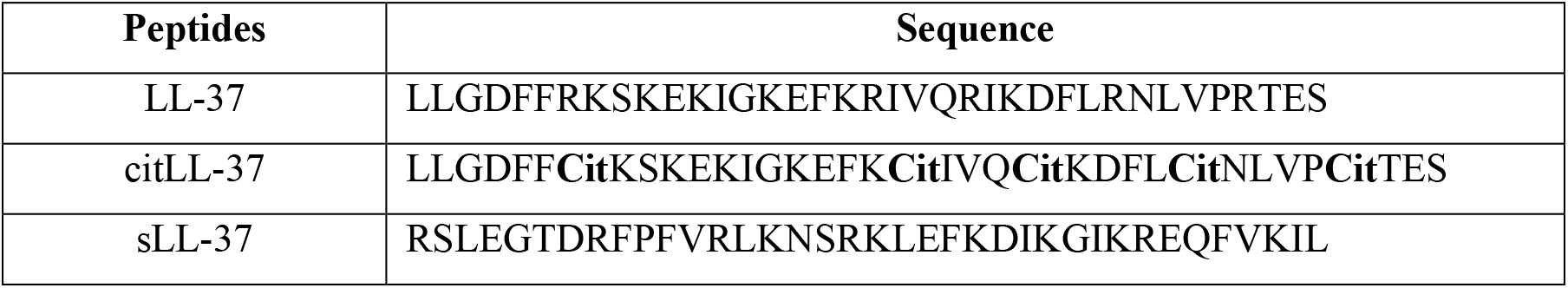
Sequences for LL-37, citLL-37, and sLL-37 peptides.

### Human bronchial epithelial cell culture

The human bronchial epithelial cell line HBEC3-KT (CRL-4051™) was purchased from American Type Culture Collection, cultured in airway epithelial cell basal medium (ATCC® PCS-300-030™) supplemented with bronchial epithelial cell growth kit (ATCC® PCS-300-040™), according to the manufacturer’s instructions and as previously described by us [32]. The cell culture medium was replaced with airway epithelial cell basal medium containing 6 mM L-glutamine, without the growth supplements, 24 h prior to stimulation with peptides.

### Targeted Transcriptomic profiling

HBEC3-KT cells were stimulated with either LL-37, citLL-37, or sLL-37 (0.50 µM each). RNA was isolated after 4 h using the MagMAX™-96 Total RNA Isolation Kit (Invitrogen; Thermo Fisher Scientific, MA, USA; Cat. No. AM1830) and the KingFisher™ Flex automated purification system (Thermo Fisher Scientific), as previously described by us [14,33]. RNA concentration was determined using a NanoDrop spectrophotometer, and RNA quality and integrity assessed using an Agilent Bioanalyzer with the Agilent RNA 6000 Nano Kit (Fisher Scientific; Cat. No. NC1783726). Total RNA (100 ng) was analyzed using the NanoString™ nCounter® Host Response Panel (Bruker Spatial Biology, WA, USA) consisting of 758 genes, including eight negative controls, six positive controls, and twelve housekeeping genes for normalization. Raw counts were normalized to positive controls and housekeeping genes using the nSolver™ Analysis Software v4.0 (NanoString Technologies) to account for technical variability. The Log2-transformed expression values were exported for bioinformatics and statistical analyses in RStudio (Version 2026.01.0+392).

PCA was performed using the prcomp function in RStudio on the Log2-normalized expression data. Delta-expression matrices were generated by subtracting the mean unstimulated control expression value for each gene prior to PCA, and the PCA results were visualized using ggplot2. Differential gene expression analyses were conducted using linear models (limma) package with empirical Bayes moderation to compare LL-37 vs unstimulated cells, citLL-37 vs unstimulated cells, sLL-37 vs unstimulated cells, and LL-37 vs citLL-37. Based on our previous studies, sLL-37 was used as the paired negative peptide. Thus, genes that showed significant changes with sLL-37 compared to unstimulated cells (Supplementary Table 1) were excluded from further analyses [9,14,34]. Volcano plots were generated using ggplot2. KEGG pathway enrichment analysis was performed using the clusterProfiler package in RStudio, with the gene universe restricted to 758 genes represented on the NanoString nCounter Host Response Panel. Genes that were significantly altered by either LL-37 or citLL-37 compared to unstimulated cells, and with significant quantitative difference in response between LL-37 and citLL-37, were analyzed using the enrichKEGG function with *Homo sapiens* as the reference organism. Enriched pathways were visualized using custom bubble plots generated in ggplot2

### Multiplex cytokine profiles

HBEC3-KT cells were stimulated with either LL-37, citLL-37, or sLL-37 (0.50 µM each). TC supernatants were collected after 24 h (n=4) and processed using the Human Cytokine/chemokine 96-plex Discovery Assay^®^ Array using the Luminex® 200™ platform (at Eve Technologies Corp, AB, Canada). Analyte concentrations flagged as out of range (OO R), indicating values below the assay’s lower limit of detection, were assigned a value equal to half of the lowest standard curve concentration for that analyte prior to Log2 transformation and statistical analysis.

### Statistical Analysis

The Log2 abundance values for genes from the NanoString analysis were analyzed in RStudio using linear models with empirical Bayes moderation. Genes with a *p*-value < 0.05 and a Log2 fold change ≥ 0.5 were considered significantly differentially expressed. PCA, volcano plots, and KEGG pathway enrichment analyses were performed using custom RStudio scripts. For the Eve Technologies Human Cytokine/Chemokine 96-Plex Discovery Assay, differential protein expression analyses were conducted using the limma package with empirical Bayes moderation. Proteins with a *p*-value < 0.05 and a Log2 fold change ≥ 0.5 were considered significantly differentially expressed.

## Acknowledgements

This work was supported by fundings obtained from the Natural Sciences and Engineering Research Council of Canada Discovery Grant (RGPIN-2026-06397). P.R. was supported by Doctoral Studentship from Research Manitoba and the Children’s Hospital Research Institute of Manitoba (CHRIM). C.L.M. was supported by studentships and awards from Research Manitoba, Mindel and Tom Olenick Award in Immunology, Asthma Canada, Canadian Allery Asthma and Immunology Foundation, Canadian Institutes of Health Research, CHRIM, and Women’s Health Research Foundation of Canada.

## Author Contributions

P.R. performed majority of the experiments, analysis of the data, prepared the figures and tables, and wrote the original manuscript. D.H.D. performed the NanoString™ nCounter® Host Response experiment and extraction of the raw dataset. M.H. provided intellectual input for optimization of the experiments and conceptualization of the study. C.L.M. performed data analysis for the cytokine multiplex panel data. C.D.P. and N.M. conceptualized the study, were responsible for funding acquisition, and provided resources and supervision. N.M. extensively edited the manuscript and provided overall supervision. All authors reviewed and edited the manuscript.

## Data Availability

All data supporting the findings of this study are available within the paper and its Supplementary Information.

## Competing Interest Statement

The authors declare no competing interests.

## Notes

### Competing Interest Statement

The authors have declared no competing interest.

