## Supplemental Information for "LL-37 and Citrullinated-LL-37 elicited Transcriptome and Cytokine Profiles in Human Bronchial Epithelial Cells: Citrullination dampens the inflammatory biosignature"

**Supplementary Table 1.** ***Genes significantly altered by scrambled LL-37.***

| **Gene** | **sLL-37** | |
|---|---|---|
| | **Log2 Fold-Change*** | ***p*-value** |
| *ZAP70* | -0.65 | 3.39E-04 |
| *APOBEC3G* | -0.72 | 4.57E-03 |
| *CSF2RA* | 0.82 | 5.58E-03 |
| *GZMH* | 0.61 | 5.64E-03 |
| *IL23R* | 0.62 | 6.83E-03 |
| *CD8A* | -0.95 | 7.44E-03 |
| *MARCO* | 0.79 | 1.71E-02 |
| *IFNK* | 0.69 | 2.36E-02 |
| *FCGR3A/B* | 0.94 | 2.55E-02 |
| *IL2RA* | -0.85 | 3.04E-02 |
| *EOMES* | -1.18 | 3.07E-02 |
| *CCL8* | -0.79 | 3.07E-02 |
| *CD45RA* | -0.59 | 3.26E-02 |
| *IFIT2* | 0.73 | 3.29E-02 |
| *CCL26* | 0.54 | 3.55E-02 |
| *HLA-DPB1* | 1.45 | 3.65E-02 |
| *CD3D* | -0.69 | 3.69E-02 |
| *IL4* | -0.65 | 3.72E-02 |
| *GAB2* | -0.62 | 3.88E-02 |
| *ADORA2A* | -1.00 | 3.90E-02 |
| *ELANE* | -0.86 | 4.16E-02 |
| *F5* | 0.84 | 4.41E-02 |

*Compared to unstimulated cells

**Supplementary Table 2.** ***Differentially expressed genes in response to LL-37 and/or citLL-37.***

| Gene | LL-37 | | citLL-37 | |
|---|---|---|---|---|
| | Log2 Fold-Change* | *p*-value | Log2 Fold-Change* | *p*-value |
| *CXCL8* | 3.48 | 2.21E-08 | 1.37 | 9.80E-05 |
| *CXCL3* | 1.58 | 1.28E-06 | 0.62 | 2.42E-03 |
| *IRF1* | -1.28 | 1.60E-06 | -0.61 | 7.07E-04 |
| *CD274* | 1.18 | 1.67E-06 | 0.39 | 8.09E-03 |
| *TRIM6* | -1.07 | 2.59E-06 | -0.50 | 1.25E-03 |
| *BCL2L1* | 1.05 | 4.13E-06 | 0.70 | 1.33E-04 |
| *CXCL1* | 1.76 | 5.31E-06 | 0.85 | 1.76E-03 |
| *NFAT5* | 1.40 | 5.93E-06 | 0.93 | 1.82E-04 |
| *RB1CC1* | 0.75 | 6.95E-06 | 0.52 | 1.55E-04 |
| *IL1A* | 1.70 | 7.78E-06 | 0.53 | 2.50E-02 |
| *CSF2* | 3.08 | 1.07E-05 | 1.09 | 1.62E-02 |
| *ATP6V1B2* | 0.78 | 1.42E-05 | 0.41 | 2.20E-03 |
| *TXNIP* | -1.45 | 1.95E-05 | -0.80 | 1.96E-03 |
| *NAMPT* | 0.81 | 2.39E-05 | 0.34 | 1.10E-02 |
| *PTGS2* | 3.85 | 2.76E-05 | 1.71 | 8.79E-03 |
| *VEGFA* | 1.97 | 3.10E-05 | 0.95 | 6.14E-03 |
| *CASP4* | 0.74 | 3.92E-05 | 0.34 | 8.61E-03 |
| *MCL1* | 1.14 | 4.39E-05 | 0.50 | 1.30E-02 |
| *CBL* | 0.91 | 4.90E-05 | 0.54 | 2.48E-03 |
| *SOCS3* | 1.70 | 4.92E-05 | 0.91 | 4.71E-03 |
| *TRAF6* | 0.78 | 5.04E-05 | 0.57 | 5.77E-04 |
| *CXCL14* | -0.76 | 5.80E-05 | -0.54 | 8.07E-04 |
| *TANK* | 0.74 | 5.87E-05 | 0.41 | 3.90E-03 |
| *SLC2A3* | 1.94 | 6.20E-05 | 1.30 | 1.29E-03 |
| *ATF4* | 0.63 | 7.01E-05 | 0.23 | 4.18E-02 |
| *TNFSF10* | -0.84 | 7.71E-05 | -0.34 | 2.73E-02 |
| *HMOX1* | 2.70 | 7.89E-05 | 1.43 | 6.92E-03 |
| *IL6ST* | 0.68 | 8.56E-05 | 0.31 | 1.47E-02 |
| *IL1B* | 1.36 | 9.37E-05 | 0.60 | 1.91E-02 |
| *PELI1* | 1.31 | 9.63E-05 | 0.59 | 1.88E-02 |
| *IL1RAPL2* | 0.88 | 9.73E-05 | 0.80 | 2.20E-04 |
| *RNASEL* | -1.56 | 1.14E-04 | -0.55 | 5.44E-02 |
| *NT5E* | 0.61 | 1.16E-04 | 0.41 | 2.23E-03 |
| *PFKFB3* | 0.93 | 1.19E-04 | 0.52 | 6.78E-03 |

| | | | | |
|---|---|---|---|---|
| *RIPK2* | 0.61 | 1.22E-04 | 0.43 | 1.58E-03 |
| *RAB7A* | 0.68 | 1.29E-04 | 0.32 | 1.79E-02 |
| *ALPK1* | -0.70 | 1.35E-04 | -0.47 | 2.38E-03 |
| *LYN* | 0.52 | 1.39E-04 | 0.26 | 1.49E-02 |
| *GK* | 1.17 | 1.53E-04 | 0.73 | 4.39E-03 |
| *IL1RL1* | 2.14 | 1.68E-04 | 1.07 | 1.53E-02 |
| *MYC* | 1.32 | 1.71E-04 | 0.92 | 2.26E-03 |
| *NEU1* | 0.82 | 1.90E-04 | 0.41 | 1.68E-02 |
| *IL20RB* | -0.84 | 2.05E-04 | -0.56 | 3.61E-03 |
| *FAS* | 0.53 | 2.30E-04 | 0.26 | 2.09E-02 |
| *MAP2K3* | 0.63 | 2.48E-04 | 0.26 | 4.60E-02 |
| *IL10RB* | -0.62 | 3.16E-04 | -0.47 | 2.06E-03 |
| *IL17RA* | -0.68 | 3.19E-04 | -0.23 | 9.31E-02 |
| *TNFRSF10B* | 0.64 | 3.53E-04 | 0.39 | 9.26E-03 |
| *CXCL2* | 1.28 | 3.69E-04 | 0.23 | 3.57E-01 |
| *CRK* | 0.61 | 3.83E-04 | 0.36 | 1.10E-02 |
| *AP1S2* | 0.54 | 3.87E-04 | 0.31 | 1.39E-02 |
| *TAB1* | -0.65 | 4.15E-04 | -0.18 | 1.90E-01 |
| *CEBPB* | 0.62 | 5.23E-04 | 0.26 | 6.25E-02 |
| *NFE2L2* | 0.52 | 6.07E-04 | 0.29 | 1.94E-02 |
| *IL24* | 1.53 | 6.20E-04 | 0.36 | 2.74E-01 |
| *MT2A* | 0.62 | 6.20E-04 | 0.38 | 1.25E-02 |
| *KDM6B* | 0.96 | 6.48E-04 | 0.38 | 8.30E-02 |
| *DTX3L* | -0.67 | 6.76E-04 | -0.43 | 1.12E-02 |
| *PLAUR* | 1.31 | 6.81E-04 | 0.62 | 4.49E-02 |
| *RNF135* | -0.91 | 7.16E-04 | -0.28 | 1.70E-01 |
| *TNF* | -0.99 | 7.26E-04 | -0.47 | 4.60E-02 |
| *ETS1* | 0.78 | 7.66E-04 | 0.46 | 1.87E-02 |
| *CASP8* | -0.54 | 8.68E-04 | -0.34 | 1.52E-02 |
| *EGLN1* | 0.66 | 8.89E-04 | 0.43 | 1.20E-02 |
| *IL1RAP* | 0.66 | 9.30E-04 | 0.45 | 9.93E-03 |
| *FCGRT* | 0.84 | 9.59E-04 | 0.57 | 1.02E-02 |
| *NAE1* | -0.70 | 1.34E-03 | -0.40 | 3.04E-02 |
| *BDKRB2* | -0.72 | 1.59E-03 | -0.47 | 1.80E-02 |
| *PIK3R3* | -0.83 | 1.66E-03 | -0.64 | 8.41E-03 |
| *PLAU* | 0.75 | 1.81E-03 | 0.40 | 4.68E-02 |
| *STAT2* | -0.51 | 2.11E-03 | -0.55 | 1.17E-03 |
| *IKBKE* | -0.63 | 2.11E-03 | -0.40 | 2.59E-02 |
| *APOL6* | -0.70 | 2.34E-03 | -0.51 | 1.44E-02 |

| | | | | |
|---|---|---|---|---|
| *IRAK4* | -0.62 | 2.44E-03 | -0.29 | 8.73E-02 |
| *NFATC3* | -0.65 | 2.76E-03 | -0.06 | 7.23E-01 |
| *SYK* | -0.56 | 2.79E-03 | -0.38 | 2.17E-02 |
| *MAVS* | -0.57 | 3.14E-03 | -0.08 | 6.00E-01 |
| *PARP9* | -0.66 | 3.26E-03 | -0.47 | 2.16E-02 |
| *ACKR3* | -0.57 | 3.40E-03 | -0.35 | 3.84E-02 |
| *MAPK8* | 0.59 | 3.63E-03 | 0.41 | 2.56E-02 |
| *GADD45B* | 0.51 | 3.89E-03 | 0.18 | 2.27E-01 |
| *PTPN4* | -0.58 | 4.06E-03 | -0.29 | 9.62E-02 |
| *IL36G* | 1.03 | 4.86E-03 | 0.61 | 5.80E-02 |
| *IFIH1* | -0.66 | 4.91E-03 | -0.35 | 8.40E-02 |
| *CXCR5* | 0.51 | 4.97E-03 | 0.51 | 4.97E-03 |
| *PRDM1* | 0.55 | 5.74E-03 | 0.16 | 3.22E-01 |
| *CYSTM1* | 0.81 | 6.47E-03 | 0.44 | 8.85E-02 |
| *PLIN4* | -0.91 | 7.64E-03 | -0.57 | 6.25E-02 |
| *IL1R2* | 0.59 | 8.63E-03 | 0.73 | 2.26E-03 |
| *IL17C* | -1.11 | 9.46E-03 | 0.51 | 1.70E-01 |
| *FCRL2* | 0.69 | 9.57E-03 | 0.62 | 1.66E-02 |
| *TLR4* | 0.64 | 1.18E-02 | 0.35 | 1.19E-01 |
| *IL11* | 0.76 | 1.35E-02 | 0.28 | 3.00E-01 |
| *IL21* | 1.13 | 1.54E-02 | 0.77 | 7.37E-02 |
| *PNOC* | 0.64 | 1.61E-02 | 0.25 | 2.90E-01 |
| *IL23A* | 0.51 | 1.62E-02 | 0.22 | 2.48E-01 |
| *IL17F* | 0.86 | 1.73E-02 | 1.30 | 1.60E-03 |
| *CD2* | 0.75 | 2.22E-02 | 0.41 | 1.70E-01 |
| *CD3G* | 0.64 | 2.32E-02 | 0.54 | 4.95E-02 |
| *IL6R* | 0.81 | 2.43E-02 | 0.36 | 2.63E-01 |
| *IL1RL2* | -0.58 | 2.47E-02 | -0.28 | 2.25E-01 |
| *CASP5* | 1.11 | 2.52E-02 | 0.59 | 1.88E-01 |
| *IFNA4/7/10/17/21* | 0.90 | 2.61E-02 | 0.89 | 2.75E-02 |
| *IFIT3* | -0.70 | 2.84E-02 | -0.43 | 1.44E-01 |
| *HLA-DPA1* | 0.67 | 2.89E-02 | 0.63 | 3.60E-02 |
| *CCRL2* | 0.86 | 2.94E-02 | 0.74 | 5.37E-02 |
| *HLA-DMB* | 0.89 | 3.00E-02 | 0.66 | 8.98E-02 |
| *OASL* | -0.59 | 3.24E-02 | -0.06 | 8.15E-01 |
| *DDIT3* | -0.51 | 3.39E-02 | -0.66 | 1.02E-02 |
| *TMPRSS2* | 0.86 | 3.55E-02 | 0.39 | 2.92E-01 |
| *SOCS1* | -0.67 | 3.64E-02 | -0.46 | 1.31E-01 |
| *HCK* | 0.76 | 3.77E-02 | 0.73 | 4.43E-02 |

| | | | | |
|---|---|---|---|---|
| *IL5RA* | 0.64 | 3.97E-02 | 0.23 | 4.17E-01 |
| *CXCL11* | 0.69 | 4.09E-02 | 1.02 | 5.97E-03 |
| *GUCY1A1* | 0.66 | 4.14E-02 | 0.66 | 4.14E-02 |
| *MAP3K5* | -0.50 | 4.17E-02 | -0.53 | 3.38E-02 |
| *SELL* | -1.07 | 4.22E-02 | -0.07 | 8.82E-01 |
| *GCA* | -0.50 | 4.36E-02 | -0.52 | 3.83E-02 |
| *LILRB2* | 0.46 | 4.72E-02 | 0.65 | 9.60E-03 |
| *HERC5* | 0.50 | 5.62E-02 | 0.71 | 1.16E-02 |
| *CCR8* | 0.76 | 6.59E-02 | 1.21 | 7.94E-03 |
| *RASGRP4* | 0.37 | 7.02E-02 | 0.71 | 3.15E-03 |
| *ACKR4* | 0.55 | 7.17E-02 | 0.98 | 4.91E-03 |
| *CCR7* | -0.93 | 7.24E-02 | -1.30 | 1.81E-02 |
| *IFNA6* | 1.28 | 8.53E-02 | 1.71 | 2.83E-02 |
| *LIF* | -0.37 | 9.22E-02 | -0.60 | 1.20E-02 |
| *GZMA* | 0.56 | 9.33E-02 | 1.03 | 6.81E-03 |
| *IL22RA1* | -0.49 | 9.60E-02 | -0.65 | 3.59E-02 |
| *CCL3/L1/L3* | 0.83 | 9.68E-02 | 1.23 | 2.18E-02 |
| *KIR2DL1* | 0.78 | 1.05E-01 | 1.09 | 3.24E-02 |
| *FAM30A* | 0.40 | 1.68E-01 | 0.62 | 4.31E-02 |
| *TNFRSF17* | 0.45 | 1.86E-01 | 0.95 | 1.35E-02 |
| *IL13* | -0.26 | 1.87E-01 | -0.55 | 1.32E-02 |
| *JAML* | 0.34 | 1.90E-01 | 0.61 | 3.16E-02 |
| *CCL17* | 0.27 | 2.56E-01 | 0.52 | 3.97E-02 |
| *PTPRC* | 0.38 | 2.85E-01 | 0.84 | 3.01E-02 |
| *IFNA2* | 0.39 | 3.12E-01 | 0.86 | 4.10E-02 |
| *OSM* | -0.17 | 5.22E-01 | -0.70 | 1.94E-02 |
| *KIR3DL1/2* | -0.15 | 6.40E-01 | 0.86 | 2.26E-02 |
| *FOS* | 0.09 | 6.97E-01 | -0.80 | 6.49E-03 |
| *IFNA14/16* | 0.13 | 7.26E-01 | 0.83 | 4.50E-02 |
| *RSAD2* | -0.08 | 7.52E-01 | 0.55 | 4.99E-02 |
| *AIF1* | -0.02 | 9.35E-01 | 0.78 | 2.95E-03 |

**Compared to unstimulated cells.*
*Grey denotes genes with Log2 fold-change < 0.5 and/or p-value >0.05 (not significant).*

**Supplementary Table 3.** ***LL-37, cit-LL-37 and sLL-37 mediated cytokine profile.***

| Protein | LL-37 | | citLL-37 | | sLL-37 | |
|---|---|---|---|---|---|---|
| | Log2 Fold Change* | *p*-value | Log2 Fold Change* | *p*-value | Log2 Fold Change* | *p*-value |
| GM-CSF | 4.33 | 3.50E-10 | 0.27 | 3.03E-01 | OOR | ns |
| MIP-3α | 2.62 | 2.13E-09 | 0.39 | 4.25E-02 | 0.51 | 1.19E-02 |
| I-TAC | 0.65 | 2.76E-07 | -0.02 | 7.46E-01 | -0.08 | 2.48E-01 |
| IL-6 | 1.02 | 2.09E-06 | 0.25 | 6.53E-02 | 0.01 | 9.19E-01 |
| IL-8 | 4.27 | 4.24E-06 | 1.47 | 2.08E-02 | 0.43 | 4.53E-01 |
| sFas | 0.38 | 5.67E-06 | 0.25 | 3.52E-04 | 0.19 | 3.07E-03 |
| IP-10 | 2.89 | 7.35E-06 | 0.93 | 3.68E-02 | 0.33 | 4.25E-01 |
| Fractalkine | 2.87 | 1.66E-04 | 0.20 | 7.18E-01 | 0.80 | 1.65E-01 |
| GROα | 2.35 | 3.95E-04 | 1.57 | 7.22E-03 | 0.90 | 9.13E-02 |
| TNFα | 1.25 | 6.45E-04 | 0.72 | 2.32E-02 | 0.27 | 3.42E-01 |
| Eotaxin | 1.25 | 1.18E-03 | 0.46 | 1.53E-01 | -0.08 | 8.00E-01 |
| sCD40L | 2.73 | 1.89E-03 | 1.08 | 1.47E-01 | 1.61 | 3.88E-02 |
| IL-18 | 0.25 | 2.43E-03 | 0.27 | 1.06E-03 | -0.05 | 4.29E-01 |
| IL-1α | 2.07 | 3.03E-03 | 0.85 | 1.57E-01 | -0.76 | 2.05E-01 |
| IFNγ | 0.68 | 4.96E-03 | 0.20 | 3.38E-01 | 0.06 | 7.62E-01 |
| RANTES | 0.65 | 6.03E-03 | 0.55 | 1.68E-02 | 0.21 | 3.00E-01 |
| GCP-2 | 0.06 | 7.38E-03 | 0.06 | 7.45E-03 | 0.04 | 4.27E-02 |
| IL-11 | 0.68 | 9.50E-03 | 1.05 | 4.80E-04 | 0.99 | 7.23E-04 |
| G-CSF | 0.80 | 2.04E-02 | 0.22 | 4.73E-01 | 0.26 | 4.04E-01 |
| HMGB1 | 0.23 | 2.18E-02 | 0.06 | 4.72E-01 | -0.01 | 9.45E-01 |
| MIG/CXCL9 | -0.30 | 2.70E-02 | OOR | ns | OOR | ns |
| TNFβ | 1.14 | 3.57E-02 | 0.41 | 4.11E-01 | -0.18 | 7.11E-01 |
| SCF | -1.46 | 3.58E-02 | 1.38 | 4.55E-02 | 1.33 | 5.24E-02 |
| IL-4 | 0.41 | 4.04E-02 | 0.15 | 4.11E-01 | -0.03 | 8.73E-01 |
| BAFF | -0.25 | 4.12E-02 | 0.00 | 9.90E-01 | -0.20 | 9.02E-02 |
| Eotaxin-2 | 0.33 | 4.68E-02 | 0.20 | 1.92E-01 | 0.25 | 1.18E-01 |
| LIF | -0.20 | 4.98E-02 | 0.01 | 9.56E-01 | 0.23 | 2.60E-02 |
| MCP-3 | 0.64 | 7.32E-02 | 0.05 | 8.73E-01 | -0.23 | 5.06E-01 |
| MCP-1 | 0.40 | 8.24E-02 | 0.00 | 9.93E-01 | -0.21 | 3.26E-01 |
| VEGF-A | 1.14 | 8.30E-02 | 0.54 | 3.88E-01 | -0.40 | 5.20E-01 |
| TSLP | -0.34 | 9.83E-02 | 0.14 | 4.79E-01 | 0.31 | 1.34E-01 |
| APRIL | 0.10 | 1.05E-01 | 0.07 | 2.33E-01 | -0.05 | 3.67E-01 |
| TARC | -0.12 | 1.19E-01 | 0.02 | 7.81E-01 | -0.04 | 5.73E-01 |
| IL-9 | 0.19 | 1.54E-01 | -0.05 | 6.78E-01 | 0.16 | 2.24E-01 |

| | | | | | | |
|---|---|---|---|---|---|---|
| CXCL16 | 0.40 | 1.55E-01 | 0.95 | 3.74E-03 | 1.05 | 1.90E-03 |
| IL-17E/IL-25 | -0.50 | 1.72E-01 | OOR | ns | OOR | ns |
| IL-13 | 0.94 | 1.74E-01 | 0.28 | 6.75E-01 | -0.37 | 5.76E-01 |
| sCD137 | 0.14 | 1.74E-01 | 0.13 | 1.78E-01 | 0.19 | 6.84E-02 |
| Granzyme A | -0.08 | 2.30E-01 | 0.04 | 5.20E-01 | -0.02 | 7.05E-01 |
| FGF-2 | -0.71 | 2.37E-01 | -0.25 | 6.76E-01 | 0.50 | 4.01E-01 |
| IL-5 | 0.02 | 2.46E-01 | 0.00 | 9.97E-01 | 0.03 | 1.14E-01 |
| TRAIL | -0.08 | 2.53E-01 | -0.02 | 7.98E-01 | 0.05 | 4.60E-01 |
| FLT-3L | 0.25 | 3.02E-01 | 0.25 | 2.92E-01 | 0.08 | 7.26E-01 |
| IFNα2 | -0.56 | 3.20E-01 | -0.28 | 6.10E-01 | -0.35 | 5.23E-01 |
| I-309 | -0.03 | 3.46E-01 | 0.04 | 2.34E-01 | 0.05 | 1.60E-01 |
| MDC | 0.04 | 3.51E-01 | 0.03 | 5.72E-01 | 0.02 | 7.24E-01 |
| IL-33 | 0.52 | 3.55E-01 | 0.43 | 4.43E-01 | -0.14 | 8.01E-01 |
| MPIF-1 | -0.03 | 3.79E-01 | 0.04 | 2.18E-01 | 0.03 | 3.79E-01 |
| IL-16 | -0.19 | 3.84E-01 | -0.18 | 4.28E-01 | -0.32 | 1.64E-01 |
| IL-1β | -0.40 | 4.06E-01 | -0.36 | 4.60E-01 | 0.22 | 6.50E-01 |
| IL-17A | 0.11 | 4.11E-01 | -0.03 | 7.99E-01 | 0.08 | 5.43E-01 |
| IL-23 | 0.09 | 4.34E-01 | 0.26 | 3.16E-02 | 0.24 | 3.91E-02 |
| IL-29 | -0.14 | 4.48E-01 | -0.19 | 3.09E-01 | -0.20 | 2.88E-01 |
| IFNω | -0.10 | 4.62E-01 | 0.10 | 4.32E-01 | 0.13 | 3.33E-01 |
| M-CSF | -0.30 | 4.79E-01 | -0.04 | 9.25E-01 | -0.36 | 3.93E-01 |
| Eotaxin-3 | -0.05 | 4.86E-01 | -0.13 | 7.02E-02 | -0.11 | 1.29E-01 |
| MIP-3β | 0.04 | 4.92E-01 | 0.08 | 1.39E-01 | 0.02 | 7.70E-01 |
| IL-10 | -0.04 | 5.07E-01 | 0.01 | 8.11E-01 | -0.01 | 7.96E-01 |
| IL-24 | -0.14 | 5.35E-01 | -0.11 | 6.21E-01 | 0.16 | 4.80E-01 |
| MCP-4 | 0.10 | 5.37E-01 | 0.32 | 5.44E-02 | 0.07 | 6.56E-01 |
| ENA-78 | -0.05 | 5.73E-01 | 0.08 | 4.28E-01 | -0.08 | 4.14E-01 |
| CTACK | 0.02 | 5.75E-01 | -0.02 | 6.83E-01 | 0.00 | 1.00E+00 |
| TGFα | -0.24 | 6.13E-01 | -0.03 | 9.53E-01 | -0.16 | 7.25E-01 |
| IL-31 | 0.09 | 6.17E-01 | 0.19 | 3.10E-01 | -0.08 | 6.63E-01 |
| IL-35 | -0.13 | 6.23E-01 | 0.09 | 7.41E-01 | -0.36 | 1.86E-01 |
| SDF-1 | 0.08 | 6.35E-01 | 0.02 | 8.90E-01 | -0.08 | 6.28E-01 |
| IL-1RA | -0.07 | 6.45E-01 | -0.40 | 1.90E-02 | -0.36 | 3.11E-02 |
| BCA-1 | -0.02 | 6.89E-01 | -0.04 | 5.68E-01 | -0.11 | 1.05E-01 |
| IL-15 | -0.09 | 6.96E-01 | 0.07 | 7.86E-01 | -0.25 | 3.11E-01 |
| IL-34 | -0.16 | 7.10E-01 | -0.11 | 7.95E-01 | 0.39 | 3.71E-01 |
| PDGF-AA | 0.18 | 7.30E-01 | 0.40 | 4.54E-01 | -0.27 | 6.07E-01 |
| MCP-2 | 0.00 | 7.53E-01 | 0.01 | 2.27E-01 | 0.00 | 7.53E-01 |
| EGF | 0.13 | 7.60E-01 | 0.09 | 8.20E-01 | -0.44 | 3.03E-01 |

| | | | | | | |
|---|---|---|---|---|---|---|
| MIP 1δ | -0.10 | 7.99E-01 | 0.00 | 9.96E-01 | -0.02 | 9.51E-01 |
| IL-20 | -0.10 | 8.00E-01 | 1.33 | 5.66E-03 | 1.71 | 9.62E-04 |
| IL-21 | 0.07 | 8.00E-01 | -0.01 | 9.81E-01 | -0.23 | 4.31E-01 |
| MIP-1α | 0.18 | 8.07E-01 | 0.14 | 8.50E-01 | -0.95 | 2.11E-01 |
| IL-28A | -0.09 | 8.44E-01 | -0.21 | 6.46E-01 | -0.07 | 8.84E-01 |
| Lymphotactin | -0.14 | 8.57E-01 | 0.48 | 5.46E-01 | -0.63 | 4.36E-01 |
| IL-12p40 | 0.06 | 8.90E-01 | 0.14 | 7.64E-01 | -0.09 | 8.41E-01 |
| IL-27 | -0.05 | 9.40E-01 | -0.73 | 3.09E-01 | -0.73 | 3.09E-01 |
| IFNβ | -0.01 | 9.43E-01 | 0.01 | 9.23E-01 | 0.00 | 9.89E-01 |
| IL-17F | -0.04 | 9.52E-01 | -0.45 | 5.15E-01 | -0.09 | 8.93E-01 |
| TPO | 0.02 | 9.57E-01 | -0.38 | 3.31E-01 | 0.35 | 3.67E-01 |
| 6CKine | -0.01 | 9.69E-01 | -0.02 | 9.07E-01 | -0.08 | 5.68E-01 |
| Granzyme B | 0.00 | 9.79E-01 | 0.07 | 5.04E-01 | 0.13 | 2.41E-01 |
| IL-22 | OOR | ns | OOR | ns | OOR | ns |
| PDGF-AB/BB | OOR | ns | OOR | ns | OOR | ns |
| sFasL | OOR | ns | OOR | ns | OOR | ns |
| IL-12p70 | OOR | ns | OOR | ns | OOR | ns |
| CCL28 | OOR | ns | -0.13 | 1.74E-01 | OOR | ns |
| Perforin | OOR | ns | 0.37 | 7.84E-02 | 0.30 | 1.50E-01 |
| IL-2 | OOR | ns | OOR | ns | OOR | ns |
| IL-7 | OOR | ns | OOR | ns | OOR | ns |
| MIP-1β | OOR | ns | OOR | ns | OOR | ns |
| IL-3 | OOR | ns | OOR | ns | OOR | ns |

**Compared to unstimulated cells. OOR=below the detectable range. Grey=non-significant changes.*

**Supplementary Table 4.** ***Concordance between transcriptomic and cytokine profiling data for LL-37- and citLL-37-responsive genes/proteins.***

| **Gene (Protein)** | **LL-37** | | **citLL-37** | |
|---|---|---|---|---|
| | **Gene Expression Log2 Fold Change (*p-value*)** | **Protein Abundance Log2 Fold Change (*p-value*)** | **Gene Expression Log2 Fold Change (*p*-value)** | **Protein Abundance Log2 Fold Change (*p*-value)** |
| *CXCL8* (IL-8) | 3.48 (2.21E-08) | 4.27 (4.24E-06) | 1.37 (9.80E-05) | 1.47 (2.08E-02) |
| *CXCL1* (GROα) | 1.76 (5.31E-06) | 2.35 (3.95E-04) | 0.85 (1.76E-03) | 1.57 (7.22E-03) |
| *IL1A* (IL-1α) | 1.70 (7.78E-06) | 2.07 (3.03E-03) | 0.53 (2.50E-02) | 0.85 (1.57E-01) |
| *CSF2* (GM-CSF) | 3.08 (1.07E-05) | 4.33 (3.50E-10) | 1.09 (1.62E-02) | 0.27 (3.03E-01) |
| *VEGFA* (VEGF-A) | 1.97 (3.10E-05) | 1.14 (8.30E-02) | 0.95 (6.14E-03) | 0.54 (3.88E-01) |
| *TNFSF10* (TRAIL) | -0.84 (3.10E-05) | -0.08 (2.53E-01) | -0.34 (2.73E-02) | -0.02 (7.98E-01) |
| *IL1B* (IL-1β) | 1.36 (9.37E-05) | -0.40 (4.06E-01) | 0.60 (1.91E-02) | -0.36 (4.60E-01) |
| *IL24* (IL-24) | 1.53 (6.20E-04) | -0.14 (5.35E-01) | 0.36 (2.74E-01) | -0.11 (6.21E-01) |
| *TNF* (TNFα) | -0.99 (7.26E-04) | 1.25 (6.45E-04) | -0.47 (4.60E-02) | 0.72 (2.32E-02) |
| *IL21* (IL-21) | 1.13 (1.54E-02) | 0.07 (8.00E-01) | 0.77 (7.37E-02) | -0.01 (9.81E-01) |
| *IL23A* (IL-23) | 0.51 (1.62E-02) | 0.09 (4.34E-01) | 0.22 (2.48E-01) | 0.26 (3.16E-02) |
| *IL17F* (IL-17F) | 0.86 (1.73E-02) | -0.04 (9.52E-01) | 1.30 (1.60E-03) | -0.45 (5.15E-01) |
| *CXCL11* (I-TAC) | 0.69 (4.09E-02) | 0.65 (2.76E-07) | 1.02 (5.97E-03) | -0.02 (7.46E-01) |
| *LIF* (LIF) | -0.37 (9.22E-02) | -0.20 (4.98E-02) | -0.60 (1.20E-02) | 0.01 (9.56E-01) |
| *IL13* (IL-13) | -0.26 (1.87E-01) | 0.94 (1.74E-01) | -0.55 (1.32E-02) | 0.28(6.75E-01) |
| *CCL17* (TARC) | 0.27 (2.56E-01) | -0.12 (1.19E-01) | 0.52 (3.97E-02) | 0.02 (7.81E-01) |
| *IFNA2* (IFNα) | 0.39 (3.12E-01) | -0.56 (3.20E-01) | 0.86 (4.10E-02) | -0.28 (6.10E-01) |
